# Thermo-optoplasmonic single-molecule sensing on optical microcavities

**DOI:** 10.1101/2023.12.13.571444

**Authors:** Nikita A. Toropov, Matthew C. Houghton, Deshui Yu, Frank Vollmer

## Abstract

Whispering-gallery-mode (WGM) resonators are powerful instruments for single-molecule sensing in biological and biochemical investigations. WGM sensors leveraged by plasmonic nanostructures, known as optoplasmonic sensors, provide unprecedented sensitivity down to single atomic ions. In this article, we describe that the response of optoplasmonic sensors upon the attachment of single protein molecules strongly depends on the intensity of WGM. At low intensity, protein binding causes red shifts of WGM resonance wavelengths, known as the reactive sensing mechanism. By contrast, blue shifts are obtained at high intensities, which we explain as thermo-optoplasmonic (TOP) sensing, where molecules transform absorbed WGM radiation into heat. To support our conclusions, we experimentally investigated seven molecules and complexes; we observed blue shifts for dye molecules, amino acids and anomalous absorption of enzymes in the near-infrared spectral region. As an example of application, we propose a physical model of TOP sensing that can be used for the development of single-molecule absorption spectrometers.

**Significance Statement:** A notable contribution to the optical detection of single molecules has been brought about by using optical microcavities, specifically whispering-gallery-mode resonators that combine optical and plasmon resonances for the most sensitive single-molecule and ion detection. Adding absorption measurements to a single-molecule technique that operates in an aqueous solution would provide powerful new detection capabilities for sensing the properties of single molecules. We demonstrate here, for the first time, the detection of absorption cross-sections of protein molecules on optoplasmonic microcavities and validate the technique with seven different molecules and complexes. A proof-of-concept for single-molecule optoplasmonic absorption spectrometers is presented, based on a thermo-optoplasmonic sensing mechanism.

## Introduction

Photonic sensing of single molecules is becoming a well-established scientific direction providing powerful instruments for biological and medical sciences. Optical techniques have made it possible to experimentally observe and manipulate single nanoparticles, molecules and atomic ions (1-7). Examples of different physical mechanisms for single-molecule studies include single-molecule imaging by optical absorption (5,6), photothermal detection scheme (8), plasmon-based photothermal spectroscopy (9), light scattering-based techniques (10), manipulations with optical tweezers (1), fluorescent microscopy (11) and several others. A noticeable contribution to the optical detection of single molecules was brought by whispering-gallery-mode (WGM) resonators. Such resonators have been initially described by Lord Rayleigh in 1910 (12); to date, optical microresonators are known as having unsurpassed quality-(Q) factors – up to 10^10^ (13) making them very sensitive to small environmental perturbations. Their basic sensing principles – resonant mode changes evaluation – exploit the seminal theory proposed in the 1940s by Bethe and Schwinger (14).

One of the most common shapes of optical microresonators used for biosensing is spheres, since they are relatively easy to fabricate from standard optical fibres and possess high Q-factors (4). However, their effective mode volumes are relatively large: for high-Q spherical resonators of up to 100 µm in diameter, the mode volume at near-infrared probing wavelengths reaches ~10^3^ µm^3^, which was a limiting factor for detecting molecules below the size of a monolayer (15,16). To achieve better localization of probing light, it was proposed to decorate WGM resonators with metal nanoparticles of ~10 nm, supporting localized plasmonic oscillations (17). WGM resonators coupled to plasmonic nanoparticles, known as optoplasmonic sensors, pave a new way for applications of WGM in sensing. Optoplasmonic single-molecule sensing becomes feasible due to the proportional perturbation of the optical microcavity induced by polarizable molecules like proteins, in tandem with the near-field enhancement of the plasmonic nanoparticle such as a plasmonic nanorod (2,3). Recent examples of optoplasmonic sensor applications have demonstrated the detection of molecular movements in solutions diluted to attomolar concentrations (18) and studying single-molecule thermodynamics and conformational changes of proteins (19). There are bright prospects for single-molecule studies with optoplasmonic WGM, from advancing single-molecule investigations to specific spectral fingerprinting of molecules (20). Notably, all-dielectric WGM microtoroidal resonators have already been used for single-particle photothermal absorption spectroscopy of nanoparticles (21) and single molecules (22). Despite this, the findings regarding single molecules, as reported by Armani *et*.*al*. in their publication, were subsequently subjected to additional scrutiny through theoretical calculations in refs. (23, 24).

We propose a novel approach for single-molecule detection on the optoplasmonic WGM sensor using the thermo-optical effect initiated by single molecules binding to a plasmonic nanorod. This method represents a departure from the prevailing trend of utilizing low-intensity light for sensing of single quantum objects, down to single photons (25-27). Instead, our approach demonstrates optical single-molecule sensing at comparatively higher power levels, leading to the discovery of the thermo-optoplasmonic (TOP) biosensing mechanism. Indeed, optoplasmonic sensing experiments have mostly been performed at low intensities of WGM excitations (1-100 µW). Nevertheless, it is important to highlight that optoplasmonic sensors typically exhibit a linear response to molecules binding to plasmonic nanoparticles. This indicates that the sizes, numbers, and optical properties of the objects being studied lead to proportional red (or blue) WGM resonance wavelength shifts based on their polarizability. Specifically, molecules such as DNA and protein with an excess polarizability in water induce a red shift in the resonance wavelength, known as reactive sensing mechanism (28, 29). Herein, we reveal that increased intensity of WGM leads to disproportional and sign-changed resonance wavelength shifts in optoplasmonic single molecule detection which subsequently can be used to estimate the absorption cross-section of single molecules. For this, we built an optoplasmonic sensor with WGMs excited at near-infrared wavelength and gold nanoparticles with near-infrared plasmon resonances to study single-protein attachment events. Changing the parameters of light coupling to the WGM resonator, Q factor of the sensor and exciting intensities, we achieve a high intensity of light that activates thermal hotspots when single proteins attach to the plasmonic nanoparticle, providing information about their absorption. The universality of this technique is confirmed directly via studying binding events for seven types of molecules and complexes: unadulterated proteins, Alexa Fluor™ 790 (Alexa) conjugated proteins, pure solution-based Alexa molecules, amino acid molecules and solution-based IRdye® 800CW (IRDye) molecules (see Supplementary, Table S1).

### Optoplasmonic sensing of proteins

Four protein samples were investigated at the first stage: 3-Phosphoglycerate kinase (3PGK), adenylate kinase (Adk) and 3PGK and Adk conjugated with Alexa, respectively; protein labelling and other experimental protocols are provided in Methods.

In our experiments, we recorded the attachment events for each molecule type within the single-molecule regime of optoplasmonic sensors. The confirmation of the single-molecule regime was established by examining survival plots (see Supplementary, Fig. S1) and similar to the approach described in ref. (3). The principal experimental scheme is illustrated in Fig. 1a. Spherical WGM resonators were made by melting SMF-28 optical fibres with a CO^2^-laser. To achieve high Q-factors of WGMs, radii of resonators were set to 45±8 µm. WGMs were excited with a 780 nm emission of a cw tunable diode laser via a prism coupler at a wide range of power levels (0.01-5.5 mW) and coupling efficiencies (6-45%). The chamber, containing the WGM resonator and of ca. 300 µL volume, is formed of a polydimethylsiloxane (PDMS) polymer sandwiched between the prism and a microscope cover glass slide. The laser was connected to a fast-speed data acquisition card (DAQ), which synchronizes the laser wavelength scanning (50 Hz) and a photodiode with a PC via a LabVIEW program when recording the transmission spectrum. WGM frequency shifts were tracked in a spectral range of several picometres around a resonance line, while the resonance full width at half maximum (FWHM) changes were also recorded. The system allows to achieve ~1 fm in spectral resolution and 20 ms in time resolution.

**Figure 1.**
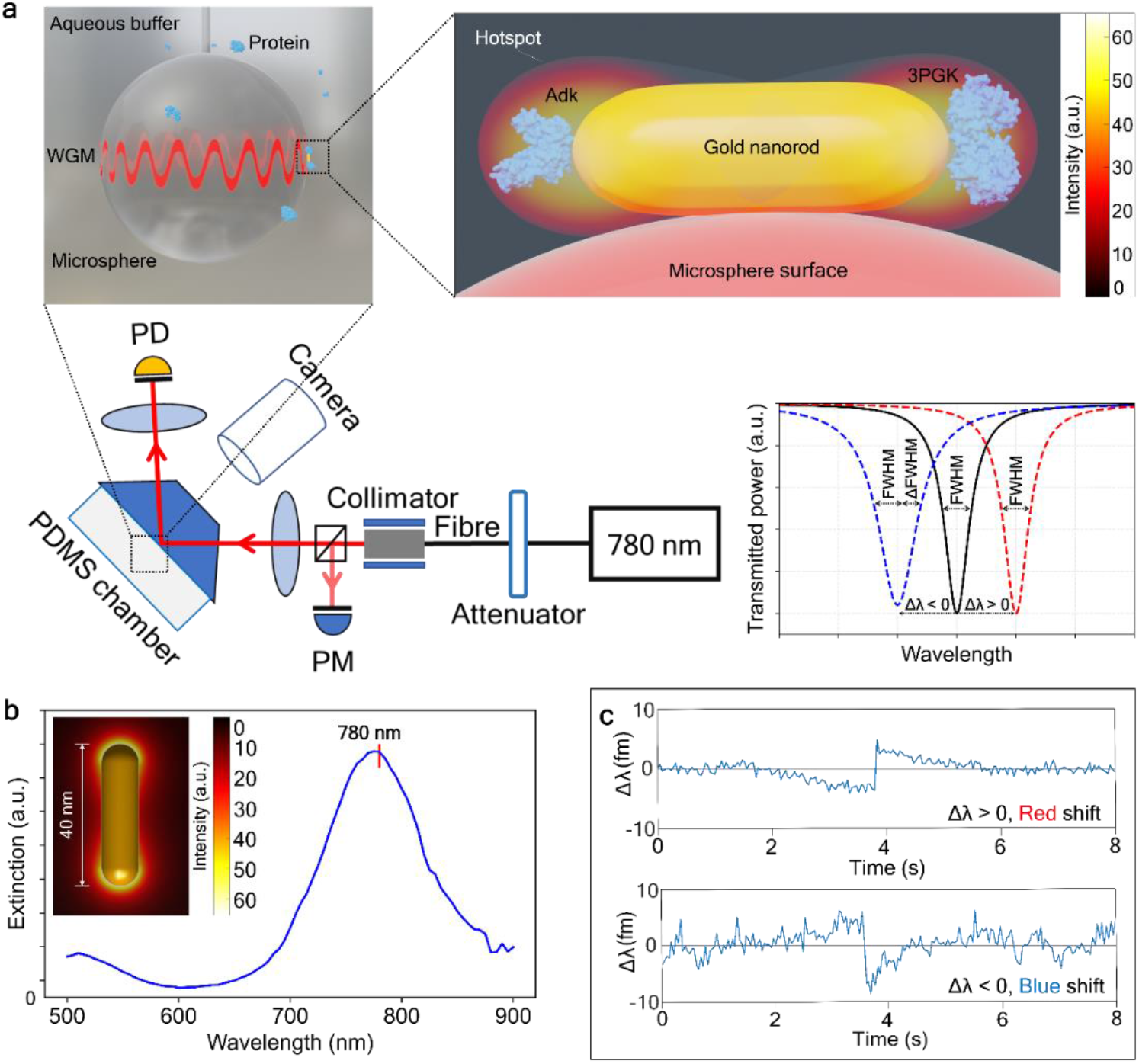
Optoplasmonic sensing. **a**, Optoplasmonic single-molecule sensor scheme. A collimated 780 nm laser beam passes through a 90/10 beam splitter: 90% – to a 50 mm focusing lens, 10% – to a power meter (PM). The incident beam (≈6°) reflects from the back side of a prism, inducing evanescent waves which excite whispering gallery modes in a ~90 µm-diameter silica microsphere placed behind the prism. The reflected beam is focused on a detector (PD); WGMs observed as dips in the transmission spectrum of the system. A PDMS chamber, containing analyte molecules, is attached to the back side of the prism. Resonance wavelength shift (Δλ) and full width at half maximum (ΔFWHM) of WGMs are tracked with the photodetector, connected to a data acquisition card (DAQ). Gold nanorods are attached to the microsphere surface. Protein samples bind to these nanorods at the tips, within an enhanced electric field that can detect perturbations of polarizability and hence the presence of protein molecules. **b**, Extinction spectrum of gold nanorods used in experiments. Inset: the electric field distribution around the nanorod with the LSPR at 780 nm. **c**, Examples of measured resonance wavelength traces showing red, *Δλ* > 0, and blue, *Δλ* < 0, wavelength shifts under the attachment of 3PGK molecules at low and high intensities of WGM, respectively.

We used an established, one step wet chemical procedure for attaching gold nanorods to the silica microsphere to assemble the optoplasmonic sensor, based on Baaske *et al*. (2) The attachments of 6-10 plasmonic gold nanoparticles, with a longitudinal LSPR (localized surface plasmon resonance) peak at 780 nm (Fig. 1b) were detected from step signals in the sensor response, mediated by a low-pH HCl solution, as a first step of the experiment. The size of the nanorods were 10 nm×38 nm, providing a negligible effect on WGM propagation; therefore, we did not observe reflected modes or mode splitting (30). Nanorod binding increases FWHM (full width at half maximum) values by roughly 100 fm (i.e. slightly reducing Q-factors) and depending on the nanorod orientation upon attachment (see Supplementary in (2)), though allows WGM sensors to be capable of detecting single molecules. In the next step of the experiment, analyte molecules (Table S1) chemically react with the attached nanorod by thiol reaction with gold, except 3PGK which utilizes a Nickel-NTA linker (see Supplementary, Fig. S2).

### 3PGK and 3PGK–Alexa

Single-molecule biosensing is based on tracking WGM resonance changes, *Δλ*, under attachment events (Fig. 1c). In our experiments, 3PGK molecules were selectively bound to the gold nanorods. Generally, under attachment events, WGM resonances are either red (*Δλ* > 0) or blue (*Δλ* < 0) shifted in relation to their initial positions, depending on the values of nanoparticle or molecular polarizability (31). However, our experiments revealed that wavelength shifts can be red and blue for the same molecules, where these shifts are mainly governed by changes of local refractive indexes. Figure 2 shows the dependence of the sign of the wavelength shift on intensity and represents an intensity-dependent diagram of single-molecule sensing with optoplasmonic sensors. For this diagram, the local evanescent intensity *I* at the tips of the nanorods (the location where the binding of single molecules can be detected, see (2,3)) was calculated by considering the effective mode volumes of TE equatorial modes, power of exciting beams, coupling values, WGM Q-factors and field enhancement around plasmonic nanorods (see Supplementary). The figure can be subdivided into three sections: reactive sensing (red shifts), near-zero shifts, and blue shifts.

**Figure 2.**
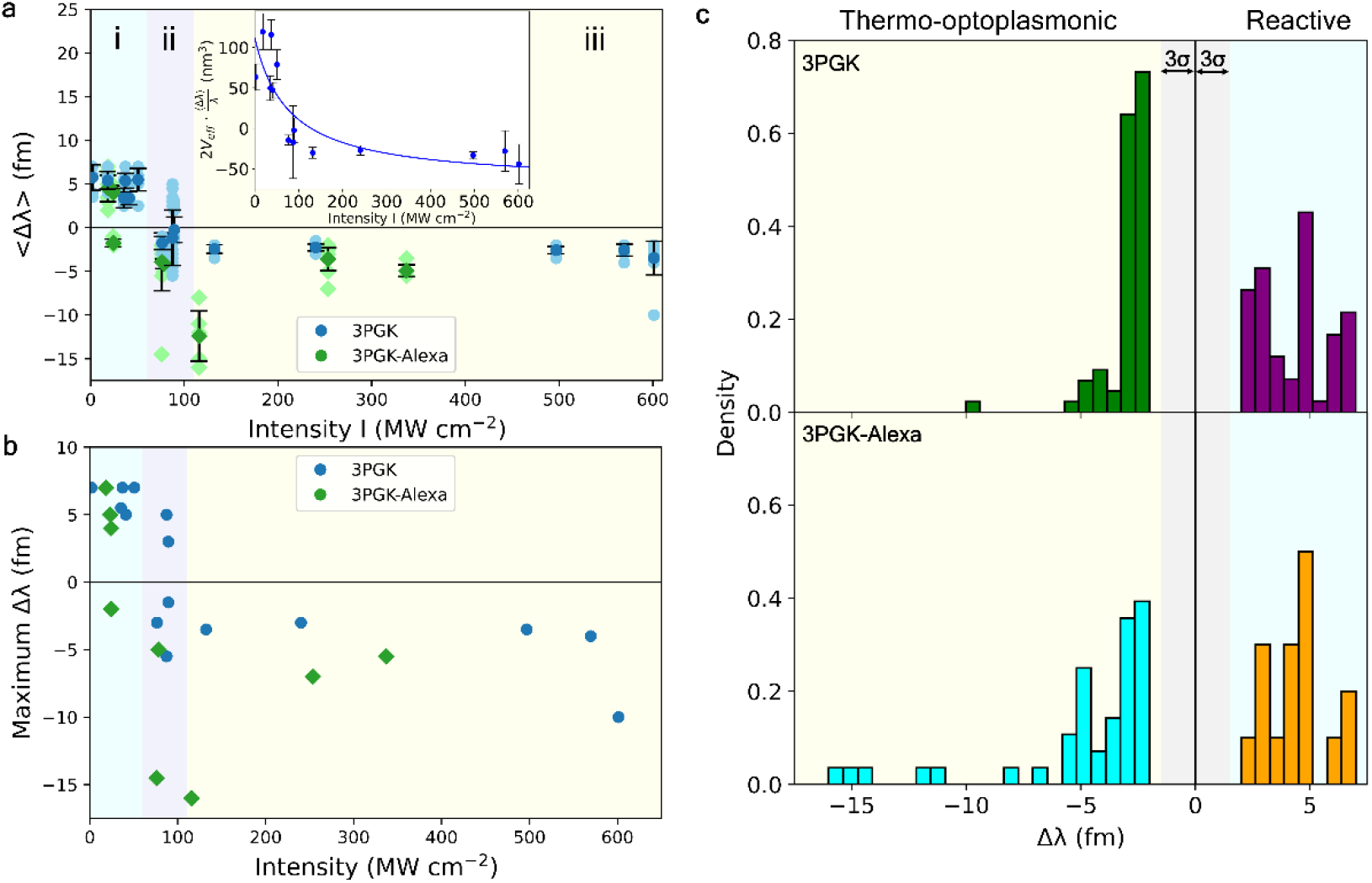
Single-molecule detection of 3PGK and 3PGK–Alexa. **a**, Optoplasmonic sensing of 3PGK (mean: dark blue circles, raw data: light blue circles) and 3PGK–Alexa (mean: dark green diamonds, raw data: light green diamonds). Averaged values ⟨*Δλ*⟩ of WGM wavelength shifts depend on the evanescent intensity I of WGM around Au nanorods. Regions (i), (ii), and (iii) represent reactive sensing, near-zero shifts, and blue shifts respectively. Inset: Dependence of 2*V*_eff_ · ⟨Δ*λ*⟩/*λ* with the effective mode volume *V*_eff_ and the wavelength *λ* = 780 nm on *I*. Symbols correspond to the experimental data while the line gives the curve fitting based on Eq. [1] – R^2^ = 0.6946. **b**, Maximal wavelength shifts vs. the evanescent intensity *I*. Maximal wavelength shifts show binding at the tips of nanorods, when nanorods longitudinal plasmonic modes are effectively excited. **c**, Histograms of wavelength shifts Δλ. 3PGK shifts grouped into the reactive sensing mechanism (red shifts – purple) and the thermo-optoplasmonic mechanism (blue shift – green). The same for 3PGK-Alexa is shown below: red shifts (orange) and blue shifts (cyan). Grey area indicates significance levels of triple the standard deviation (3*σ*).

#### (i) Reactive sensing mechanism

The first section of the diagram (Fig. 2a, i, and Fig. 2b), ending at *I*∼60 MW cm^-2^, corresponds to the conventional reactive sensing mechanism (2, 32) when binding events cause changes of the resonant wavelength – positive wavelength shifts *Δλ* > 0, as presented in Fig. 1c. During attachment events, 3PGK molecules cause a strong response, changing the polarizability in the evanescent field of the plasmonic nanorod coupled to the WGM resonator. At such intensity levels, 3PGK binding events demonstrate, independent of the evanescent intensity, wavelength shifts *Δλ* equal to 6 fm, with standard deviations of signals (*σ*) within 1 fm (Fig. 2c). Extracting signals from the WGM transmission spectra measured with optoplasmonic sensors is described in Materials. Notably, the attachment events of molecules to nanorods at low intensities of WGM do not significantly affect the spectral width (FWHM) of the resonances (Fig. 3d). Variation of the step heights of wavelength shifts seen for resonators of the same size are attributed to differences in the nanorod binding location with respect to the WGM field profile and binding orientation of the nanorod with respect to the WGM polarization (2).

**Figure 3.**
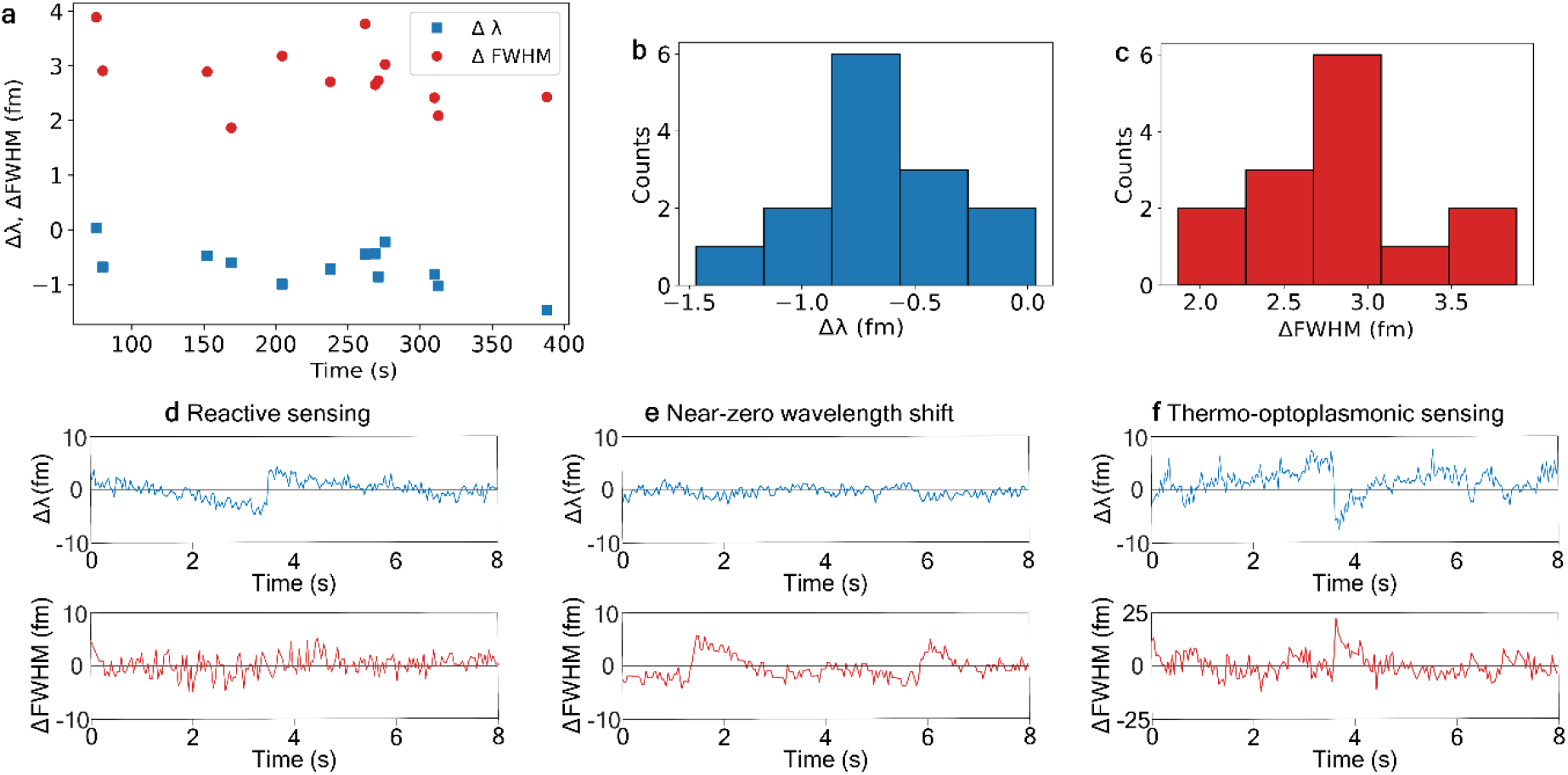
**a**, 3PGK optoplasmonic sensing at the evanescent intensity *I* = 89.3 MW cm^-2^. Near-zero resonance wavelength shifts Δ*λ* (blue squares) are accompanied by the changes of full width at half maximum (ΔFWHM) (red circles). Changes of the resonance wavelength Δ*λ* are below the noise level (i.e., triple standard deviation 3*σ*) and are considered to be effectively zero. **b**, Histogram of number of attachment events (Counts) vs wavelength changes Δ*λ*. Majority of shifts found between −0.8 and −0.6 fm, well below the noise level (3*σ* = 1.5 fm). **c**, Histogram of number of attachment events (Counts) detected via FWHM changes vs ΔFWHM. Majority found between 2.75 and 3 fm, greater than the noise level (3*σ* = 1.8 fm). **d–f**, Examples of wavelength shifts and FWHM changes. **d**, 3PGK reactive sensing with Δ*λ* > 0 and ΔFWHM ≈ 0. e, Near-zero wavelength shift with Δ*λ* ≈ 0 and ΔFWHM > 0. **f**, Thermo-optoplasmonic sensing with Δ*λ* < 0 and ΔFWHM > 0.

#### (ii) Near-zero wavelength changes

Region (ii) of Fig. 2a reflects significant changes in the mechanisms of sensing, where the wavelength shifts become near-zero (Fig. 3a, b), or protein attachment becomes effectively undetectable by resonance wavelength changes. Near-zero wavelength changes occur at specific WGM intensities where the reactive sensing (positive effect on *Δλ*) and TOP sensing (negative effect on *Δλ*) regimes balance, cancelling each other out and resulting in a net-zero effect on WGM *Δλ*. However, the single-molecule attachment events were still observable via step-like changes of FWHM (Fig. 3a, e). The FWHM of the resonances become wider by 3 fm on average (Fig. 3c), relating to increased losses of the WGM resonator. Note that the FWHM at lower intensities was measured as non-responsive to the binding events of 3PGK molecules (Fig. 3d). The difference of FWHM behaviour can be elucidated by considering the Q-factors of WGM resonators. These take into account several factors, including most notably: scattering, material losses, losses related to resonator radii and radiation losses (20). Attachment of 3PGK to optoplasmonic sensors does not lead to changes in the resonator’s geometry and therefore, at low intensities, does not contribute to resonator scattering and FWHM will not change during binding events (we presume that molecular scattering under attachment events could change values of FWHM but within 3*σ*). Q-factors and FWHM are not usually related to absorption of WGM energy by the molecules under test (20). However in the present case where FWHM changes occur at high WGM intensities when 3PGK binds (Fig. 3f), absorption of WGM energy is the only factor that can be considered to cause the FWHM response. This demonstrates that thermo-optical sensing can therefore be used for direct estimation of absorption by the molecules.

#### (iii) Blue shift

Effectively, the second (ii) and third (iii) regions of Fig. 2a represent the same mechanism of WGM resonances changes, (i.e. absorptive properties of molecules). The third region of the diagram (Fig. 2a, iii) represents blue WGM resonance wavelength shifts observed as binding events of 3PGK molecules to Au nanorods at higher local intensity (*I* >89.3 MW cm^-2^). The dependence of the sign of wavelength shifts on the intensity reveals a new mechanism of single-molecule detection with optoplasmonic sensors, which is related to the strong mutual influence of intensity of WGMs and molecules under test. Binding events of 3PGK molecules to the nanorod appear as blue shifts (*Δλ* < 0) accompanied by the partial absorption of optical energy by different amino acids of the 3PGK molecule (*ΔFWHM* > 0), see Fig. 3f. In normal conditions, when intensities are relatively low, 3PGK molecules have a strong absorption band centred at 276 nm, which corresponds to the absorption by the aromatic amino acids of 3PGK molecules: tryptophan, phenylalanine, histidine and tyrosine (see Supplementary, Table S2). Our results reveal that when a single 3PGK molecule is attached to a single plasmonic nanoparticle, it demonstrates absorption bands in the near infrared spectrum due to tryptophan residues. In far-field spectroscopy, tryptophan molecules have absorption bands in the UV range, around 280 nm. As it was recently reported (33, 34), tryptophan molecules demonstrate red-shifted bands both as phosphorescence and absorption. Here, we experimentally demonstrate that new optical transitions are possible under strong perturbation caused by the plasmonic nanoparticle (see details in section **Tryptamine and pure dye molecules** below).

Initiated by a 3PGK binding event, energy is absorbed by 3PGK molecule which causes a local temperature increase of the water around the binding location, the timescale of which is unresolved because of the limited time resolution of the sensor (20 ms). Local heating of water makes the local refractive index smaller (water has a negative *dn/dT*). Such localized heating of water result in blue resonance wavelength shifts. This reveals the sensing mechanism, which is caused by thermal changes initiated by the binding of molecules on higher-intensity optoplasmonic sensors, or TOP sensing. Top sensing, despite employing a distinct plasmon-enhanced mechanism and resulting in the reported negative wavelength shifts, shares similarities with the thermo-optic mechanism proposed by Armani *et al*. (22, 23). Theoretical confirmation of the single-molecule findings in the context of Armani *et al*.’s work is still ongoing (23, 24).

Including the local thermal effect, the average value ⟨*Δλ*⟩ of resonance wavelength shifts induced by single protein molecules binding within the plasmon-enhanced near field (aka plasmonic hotspot) of the optoplasmonic sensor is formulated as (see Methods)

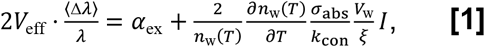

with the effective mode volume *V*_eff_ of WGM, the excess polarizability α_ex_ and absorption cross-section *σ*_abs_ of the protein, the refractive index *n*_w_(*T*) of water at the temperature *T*, the water’s thermal conductivity *k*_con_ (in units of W m^-1^ K^-1^), the effective volume of the heated water *V*_w_, and the effective heat transferring length *ξ*. It should be noted that unlike the conventional definition of the mode volume of WGM (35), *V*_eff_ here is defined based on the local light intensity at the nanorod’s hotspot (see Supplementary) and it has already included the LSPR-induced enhancement of the local electric-field intensity of the gold nanorod. The left-hand side of Eq. [1] is completely related to the microcavity (e.g., the mode volume and the resonance shift) and all environmental perturbations appear on the right-hand side of Eq. [1]. Two facts contribute to the resonance shift ⟨*Δλ*⟩ of WGM: *(i)* as usual, ⟨*Δλ*⟩ may arise from the excess polarizability α_ex_ of the protein changing the local refractive index of the WGM microcavity. (*ii)* The bound molecule absorbs the light energy, raising the local temperature and changing the refractive index of the microcavity’s surrounding medium (i.e., aqueous buffer), resulting in an extra resonance shift. Compared to the thermal effect of water, the thermo-optic effect of the microsphere is negligible because of the small rate of change of the refractive index of microsphere with respect to the temperature (see Supplementary). In the low-intensity limit *I*∼0, 2*V*_eff_ · ⟨Δ*λ*⟩/*λ* approaches α_ex_ and is positive. As *I* is enhanced, the thermal effect-induced resonance shift component grows and the positive value of 2*V*_eff_ · ⟨Δ*λ*⟩/*λ* is reduced. For a large enough *I*, 2*V*_eff_ · ⟨Δ*λ*⟩/*λ* becomes negative. In general, the effective heat transferring length *ξ* depends on the local electric-field intensity *I*. Under the linear approximation, we express *ξ* as *ξ* = *ξ*_0_ + *βI*, where the constant *ξ*_0_ approximates the radius of the protein molecule (i.e., the heat transferring distance cannot be smaller than the size of the heat source) and the parameter *β* may be derived from the curve fitting. Substituting the typical values of 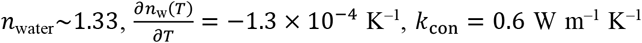 (taken from refs. (36-38)), *V*_w_ = 3.3 × 10^−23^ m^3^, and *ξ*_0_ = 3.5 nm for 3PGK, we obtain α_ex_ = 1.1 × 10^−25^ m^3^ with the 95% confidence interval (0.7 × 10^−25^, 1.6 × 10^−25^) m^3^, *σ*_abs_∼7.5 × 10^−16^ cm^2^ with the 95% confidence interval (4.2 × 10^−16^, 10.7 × 10^−16^) cm^2^, and *β* = 0.045 nm/(MW/cm^2^) with the 95% confidence interval (0.034,0.056) nm/(MW/cm^2^), from the curve fitting (Fig. 2a). The large absorption cross section for 3PGK at 780 nm is observed for the enzyme molecules attached to plasmonic nanorods, providing enhanced near fields, of optoplasmonic microcavities at sufficient optical power, suggesting that molecular transitions are excited in the TOP sensing approach that are normally weak in standard absorption spectrometry (see Supplementary, Table S3).

To confirm the mechanism explained, a series of experiments were performed for 3PGK complexes, where a dye (Alexa Fluor™ 790) with a molecular absorption peak spectrally close to the maximum of WGM wavelength, i.e. 780 nm, was selected to be covalently attached to 3PGK. The result of single-molecule 3PGK–Alexa binding is plotted in Fig. 2a and b as green dots. The results obtained were similar to experiments with unlabelled 3PGK, where positive wavelength shifts at lower intensities were obtained. They have the same values, 6 fm, when sensing by the reactive mechanism at low *I* because the structure of 3PGK and molecular weight were changed insubstantially by the Alexa label (Table S1). The WGM wavelength changes are switched to blue shifts at higher intensities of WGMs but with greater magnitude than unlabelled 3PGK. In fact, 3PGK–Alexa molecules show blue shifts at even lower intensity than non-labelled 3PGK molecules. The values of resonance shifts demonstrate big variation, likely related to the different positions of molecules on the nanorod; Fig. 2b shows maximal wavelength shifts corresponding to the position of 3PGK at the tips. In these conditions, the previously non-observable optical transitions in tryptophan have quite similar values of wavelength shifts to their counterparts in Alexa Fluor™ 790. The greater magnitude of negative shift and TOP sensing at lower *I* is due to increasing the absorption of the 3PGK molecules by conjugating Alexa, creating an additive effect to enhance the TOP mechanism.

We exclude from our aforementioned analysis the possible fluctuations of temperature due to nanorods heating effects because the proteins are added to the chamber and detected when the sensor is in the steady-state temperature regime. Nonetheless, the increased intensity of WGM causes increased background heating of nanorods (see Supplementary), that could make blue shift values upon protein binding smaller at higher intensities. We also consider that increased temperature due to WGM radiation absorption may also cause increasing temperature of the microsphere. However, using values of thermal conductivities of water, *k*_w_ = 0.6 W m^−1^ K^−1^, and silica is *k*_s_ = 1.38 W m^-1^ K^-1^, supposing that the local temperature of silica is same to the local temperature of water (i.e., the local temperature increase in silica is same to the local temperature increase in water) under the thermal equilibrium, we came to the conclusion that the change of refractive index of water with respect to the temperature is 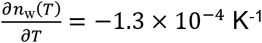. The change of refractive index of silica with respect to the temperature is 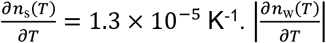 and is one order of magnitude larger than 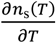. That is to say, under the same local temperature increase, the change of refractive index of water is 10 times larger than that of silica. According to this, the0020effect of the silica temperature increase on the resonance shift, which is generally a competitive process, is much smaller in comparison to the water temperature increase.

### Adk and Adk–Alexa

To provide more information about TOP sensing, we investigated another protein (Adk) and protein-dye complex (Adk-Alexa). The results are summarized in Fig. 4. In contrast to 3PGK, Adk binding causes smaller wavelength shifts of 3 fm, due to Adk molecules being almost twice smaller (44 kDa for 3PGK vs. 24 kDa for Adk). This smaller size also causes a weaker response when Adk–Alexa complexes are attached to the sensor both at high and low intensities of WGM. Indeed, the dominant mechanism of local heating is related to the absorption of Alexa molecules followed by heating the Adk–Alexa complex. Alexa-790 dye absorbs WGM radiation and transforms its energy into heat via non-radiative relaxation. The quantum yield of luminescence of Alexa in solution is about 10%, however, protein–Alexa complexes bound to the sensor have their luminescence quenched. Therefore, we expect the quantum yield to be of units of percent and hence the energy absorbed from WGM is released as heat. If we consider heat capacity values of 3PGK and Adk are roughly equal and the absorption spectra with efficiency of the Alexa–labelled subject proteins reflecting their molecular weight (see Supplementary), then the smaller mass of Adk directly indicates smaller heat absorption, i.e., blue shifts for complexes of 3PGK–Alexa and Adk–Alexa will be observed at different intensities.

**Figure 4.**
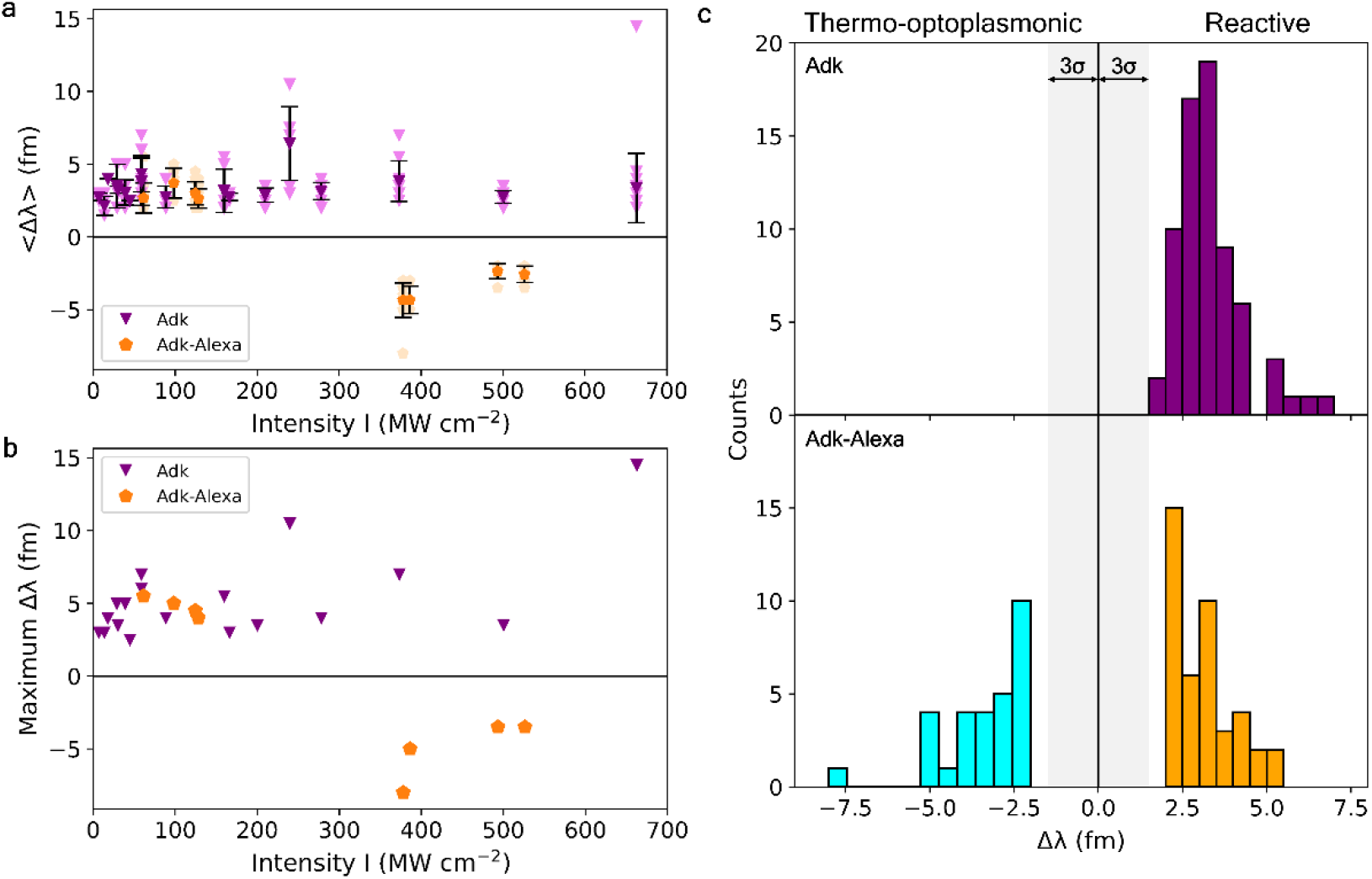
Single-molecule detection of Adk and Adk–Alexa. **a**, Optoplasmonic sensing of Adk (purple triangles) and Adk–Alexa (orange pentagons). Averaged values of WGM wavelength shifts ⟨Δ*λ*⟩ depend on the evanescent intensity *I* of WGM around Au nanorods. **b**, Maximal wavelength shifts at each intensity *I* of WGM. Maximal wavelength shifts show binding at the tips of nanorods. **c**, Histograms of wavelength shifts. Histograms of Adk (upper) binding shifts by the reactive mechanism (purple). Adk–Alexa (lower) grouped into reactive sensing (orange) and blue shifts (cyan). Grey area indicates noise levels of triple standard deviation, 3*σ*.

Note that the value of ⟨*Δλ*⟩ both for 3PGK–Alexa and Adk–Alexa complexes is close to -5 fm, also suggesting that Alexa absorption is the ruling mechanism. An important observation is that Adk molecules themselves, without Alexa conjugation, have no absorption around 780 nm and do not demonstrate blue shifts at high intensities. The crucial difference between absorption of 3PGK and Adk at 780 nm is related to the presence of tryptophan in the composition of 3PGK and absence in Adk.

### Tryptamine and pure dye molecules

In order to confirm tryptophan’s involvement in TOP sensing, it was appropriate to test the binding of small-molecule tryptophan to the surface of gold nanorods during WGM. However, competition between the amine and carboxyl group (see Supplementary, Fig. S5) for binding to the nanorods yielded spiked events rather than step-like binding events. To observe Δ*λ* in a step-like manner, a similar small molecule with the same functional group and optical properties was used: tryptamine. At pH 10, the amine group will be deprotonated and lone-pair of electrons available for interactions with the nanorod surface.

Upon binding of tryptamine to nanorods, we found a very similar trend of the Δ*λ* profile to 3PGK: intensity-dependent changes of the sign of resonant wavelength shifts upon binding. This suggests the TOP sensing effects of 3PGK are likely due to the intrinsic tryptophan residues present in 3PGK, and absence of TOP sensing in Adk due to lack of tryptophan residues. This is likely as a result of effects seen similarly in surface–enhanced resonant Raman scattering (SERRS), where electron transfer from excited plasmons in nanoparticles can form species with visible excitation bands. Sloan-Dennison *et al*. (39) demonstrate the ability of tryptophan to undergo these chemical changes when bound to plasmonic nanoparticles during Raman spectroscopy, forming a Trp-• species with absorption bands that span across the visible range, including at 780 nm. We propose the mechanism of apparent forbidden transitions in this case occurs as follows: the ≈ 780 nm WGM excites plasmons in the nanorods, resulting in electron transfer from the nanorod to indole ring of tryptophan/tryptamine. This same 780 nm WGM can excite and allow observation of optical transitions in the newly formed Trp-• species, which when relaxing to the ground state releases energy as heat, resulting in the characteristic blue shifts of TOP sensing.

Similarly, the intensity-dependence may be due to the requirement to provide sufficient energy to allow electron transfer. This may be most efficient around 70-130 MW cm^-2^ for proteins (40). Greater variability of tryptamine than larger protein molecules could also be explained due to its small-molecule nature. This is likely due to tryptamine’s closer position to the nanorod (see Fig. S2 in Supplementary). Local heating effects and evanescent intensity are more enhanced, resulting in wavelength shifts at the same apparent intensity that can show reactive and TOP sensing.

Finally, the TOP sensing mechanism was tested for attachment events of pure Alexa and IRDye molecules at different intensities. Binding of these molecules to the sensor occurred via interactions of sulfate groups with the gold nanorods (see Methods). Fig. S6 summarizes results of these control experiments. Upon Alexa molecules binding, sign-changing behaviour is observed: red-shifted resonances changes at intensities of up to 60 MW cm^-2^, followed by blue-shifted resonances at larger intensities; i.e. similar sign-changing trend to 3PGK, 3PGK-Alexa, Adk-Alexa and Tryptamine.

Binding of IRDye molecules demonstrate features of the TOP sensing mechanism. Resonant wavelength shifts are switched from red to blue at an even smaller level of intensity. The presence of negative shift signals in the dye datasets was enough to confirm our hypothesis. This demonstrates that absorption at the WGM wavelength by molecules attached to the plasmonic nanorods will allow TOP sensing effects. Near-zero averaged values of dye wavelength-shifts at high intensities are likely due to additional effects such as photodecomposition of dye molecules as well as IRDye800 and Alexa790 being small molecules and able to explore the surface roughness of gold nanorods, due to having transient/lower affinity interactions that result in both binding and unbinding signals, but also due to having multiple sulfate-mediated binding sites for the gold nanorods.

## Conclusion

We have demonstrated single molecule sensing of proteins and proteins labelled with organic dye molecules. For the first time, it was shown that WGM sensing, improved with plasmonic nanoparticles, is dependent on the intensity of WGM modes. At low intensity, sensing occurs through the reactive mechanism: red or blue shifts of WGM resonances depending on polarizability of molecules under test and their refractive index. We established that at high intensity levels, sensing can occur through a different mechanism: through the absorption of energy by molecules under test followed by heating the surrounding solution, causing blue shifts of the WGM. We have coined this mechanism thermo-optoplasmonic (TOP) sensing. The most exciting part of our results shows that by using thermo-optoplasmonic sensing, it is possible to define the absorption cross-section of single molecules due to its relation to WGM wavelength shift.

Equation [1] establishes the relation between the absorption cross-section of molecules and WGM wavelength shifts. This relation serves as a model showing that optoplasmonic sensors can be used as single molecule spectrometers. Further works can be performed via tracking multiple WGM resonances and contribute to studies of photophysical properties of an enormous number of molecules at the single molecule level.

## Materials and Methods

### Enzyme Binding to Optoplasmonic Sensor

The optoplasmonic sensor consists of two major components, the spherical silica microresonator and plasmonic gold nanorods. 6-10 cetrimonium bromide (CTAB) coated gold nanorods (Nanopartz A12-10-780-CTAB) with plasmon resonance at 780 nm were attached to WGM microspheres in 0.02 M HCl, monitored by changes in WGM resonance wavelength (Δλ) and full width at half maximum (FWHM). Poly-L-lysine-polyeythylene glycol (PLL-PEG) can be used at this stage to prevent non-specific binding to the surface of the silica microsphere, but was found to not necessary in this study.

3PGK/3PGK-Alexa immobilisation was performed by modifying the gold-nanorod surface with thiolated nitrilotriacetic acid (NTA). A mixture of 50 µM dithiobis(C2-NTA) (Dojindo D550), 450 µM Thiol-dPEG®4-acid (Sigma-Aldrich QBD10247) and 250 µM TCEP-HCl (tris(2-carboxyethyl)phosphine-HCl) was incubated for 10 mins before mixing in a 1:30 ratio with 50 mM citrate buffer and 1M NaCl and submerging the microresonator in the solution for a further 20 mins in the chamber. The chamber and resonator were washed with 50 mM HEPES. The NTA molecules, on the surface of the nanorods, are then charged with nickel ions by submerging the microresonator in 0.1 M nickel sulphate for 2 mins and chamber finally washed and filled with 50 mM 4-(2-hydroxyethyl)-1-piperazineethanesulfonic acid (HEPES). 2 µL of 0.1 mg mL–1 3PGK or 3PGK-Alexa was then added to the chamber, while monitoring the resonance shift Δλ of WGMs, relying on the Ni-NTA to His-tag interaction for enzyme immobilization onto the nanorod surface. Adk/Adk–Alexa immobilization was performed by direct covalent attachment via gold-thiol interactions, relying on a C-terminal Cys residue. To do so, the chamber was filled with 50 mM TCEP and 50 mM HEPES, and 2 µL of 0.5 mg mL–1 Adk or Adk–Alexa added to the chamber and immobilization monitored by shifts in Δλ of WGMs.

### Tryptamine and Dye Binding to Optoplasmonic Sensor

Tryptamine binding was performed via amine lone-pair interactions with gold nanorod surfaces at pH 10 in a 50 mM bicarbonate buffer. 2 µL of 1 µM tryptamine (Santa Cruz SC-206065) was added to the chamber in order to observe steps in the Δλ of the WGM.

Alexa Fluor™ 790 (ThermoFisher A30051) and IRDye® 800CW (LI-COR 929-70020) binding to the sensor was performed via interactions of sulfate groups with the gold nanorods. This was performed at pH 7.5 in 50 mM HEPES at concentrations of 13.3-26.7 nM.

### Labelling of 3PGK and Adk with Alexa Fluor™ 790

3PGK and Adk molecules were labelled with Alexa Fluor™ 790, using the Succinimidyl Ester form (ThermoFisher A30051), reacting with free amine groups on the protein surface. Alexa Fluor™ 790 was dissolved in DMSO to 10 mg mL–1 and mixed with a 3 mg/mL solution of protein in 50 mM HEPES and 0.01 M sodium bicarbonate to a final concentration of Alexa Fluor™ 790 of 0.833 mg/mL. Mixture was incubated while shaking and protected from light for 1 hour. To separate protein from free Alexa Fluor™ 790 and buffer exchanged to an appropriate buffer, size exclusion chromatography (SEC) was performed. SEC was performed using HiLoad® 16/600 Superdex® 75pg column (Cytiva 28-9893-33) with an elution buffer of 20 mM HEPES, 150 mM NaCl (pH 7.5) over 1.5 column volumes. Fractions collected and fractions selected through correlation of absorption at 280 nm and 700 nm, and confirmed by SDS-PAGE analysis. Selected fractions pooled and concentrated using Vivaspin® 20 3 kDa MWCO PES (Cytiva 28-9323-58) concentrators by centrifugation at 3 kG for 15 mins, and repeated until volume was <500 µL.

### Data processing

A graphical user interface developed in MATLAB for processing the WGM time traces was used similarly to Supplementary materials (19). A Labview program was used to record and process the WGM spectra used to track the WGM resonance position. Once, the WGM time traces are obtained, the data were analysed for peaks using the MATLAB GUI. First, drift correction caused by slow variations of temperature was applied to remove slow variations of the resonance traces. A first-order Savitzky-Golay filter with a window length depending on the sampling rate was applied to the signal. Second, step-like wavelength traces were analysed in the MATLAB program to find the resonance shift *Δλ* values corresponding to proteins binding.

Hence, the signal can be close to noises but has a higher amplitude either in wavelength or in FWHM. We quantify useful signals as all steps with amplitude higher than 3*σ* (the standard deviation of the background) of that same sample. The value of σ was evaluated by dividing the WGM time trace into windows of *N* points and evaluating the standard deviation of each *N*-point window. Typically, the value of *σ* is 0.4-0.5 fm, that increases with increased power up to three times. Diagrams (Fig. 2 and 4) combined consolidated data of 451 signals over 50 different experiments. Figure S5 consolidates 117 signals from 10 experiments.

## Acknowledgments

N.T. is grateful to Dr Simona Frustaci (University of Exeter) for preparation of 3PGK samples and Katya Zossimova (University of Exeter and University of Freiburg) for help with field distribution calculations. M.H. thanks Dr Stefan Bagby (University of Bath) for their supervision. We thank Dr Sivaraman Subramanian for useful discussions (University of Exeter) and Dr Srikanth Pedireddy (University of Exeter) for useful discussions and preparing Fig. S2 in Supplementary Materials.

