## Supplementary Materials for "Thermo-optoplasmonic single-molecule sensing on optical microcavities"

### Thermo-optoplasmonic single-molecule sensing on optical microcavities- Supplementary Material

**Table S1 | Characteristic values of tested molecules and complexes**

| Sample | 3PGK | 3PGK-Alexa | Adk | Adk-Alexa | Tryptamine | Alexa | IRDye |
|---|---|---|---|---|---|---|---|
| **Absorption peak, nm** | 276 | 779 | 271 | 777 | 296 | 780-784 | 778 |
| **Molecular weight, Da*** | 44,376 | 46,126 | 23,999 | 25,749 | 160 | 1,750 | 1,166 |
| **Extinction coefficient at 280 nm, $cm^{-1}M^{-1}$** | 22,920 | 43,700 | 10,430 | 31,200 | 5,405 | 20,784 | 7,200 |
| **Extinction coefficient at 780 nm, $cm^{-1}M^{-1}$** | ~0 | 167,000 | ~0 | 95,000 | ~0 | 260,000 | 240,000 |
| **No. of tryptophan residues (indole rings)** | 2 (2) | 2 (2) | 0 | 0 | (1) | † | – |

* As predicted by ExPASy ProtParam online tool.[1]
† Alexa Fluor™ 790 structure unpublished

**Confirmation of Single-Molecule Binding Regime**

When collecting all binding shifts where $\Delta\lambda > 3\sigma$, we find the wait time ($\Delta t$) to fit a single-exponential following $P(0, \Delta t) = e^{-R\Delta t}$ where $R$ is the rate constant of binding, as also outlined in other publications.[2] These survivor functions (Fig. S1) demonstrate binding to be on the single-molecule level. Deviations from a perfect exponential decay may be due to non-homogenous distribution of analyte in the solution due to temperature or diffusion variations.

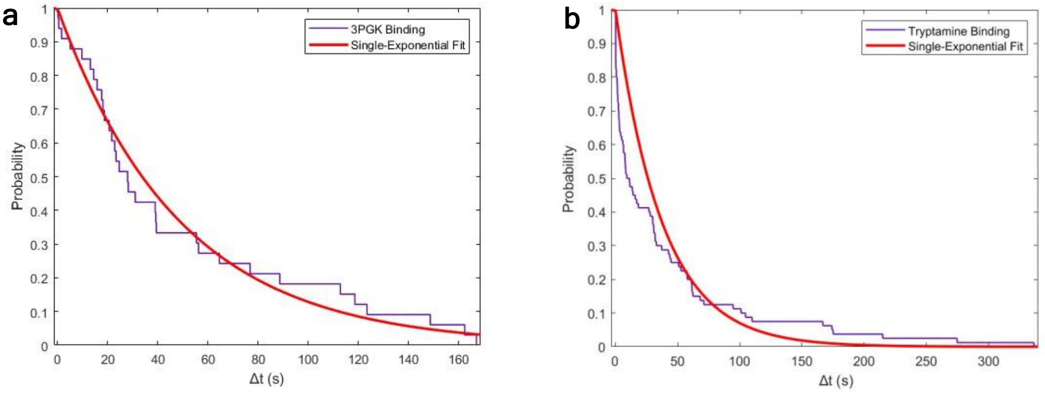


**Fig. S1. a.** Survivor function of 3PGK binding wait time ($\Delta t$) to the optoplasmonic sensor (purple) showing exponential decay where $P(0, \Delta t) = e^{-R\Delta t}$ (red). **b.** Survivor function of tryptamine binding wait time to the optoplasmonic sensor (purple) showing exponential decay (red).

##### Binding regime of all molecules

The molecules tested here bind by slightly different methods, dependent on their specific chemistries. 3PGK proteins bind via a His-tag-Ni-NTA linkage, Adk via direct cysteine sulphur-gold bonding, tryptamine via amine-gold interactions and IRDye800Cw and Alexa790 via sulfate-gold interactions (Fig. S2). These binding regimes are specific and non-random, ensuring they bind in a single orientation.

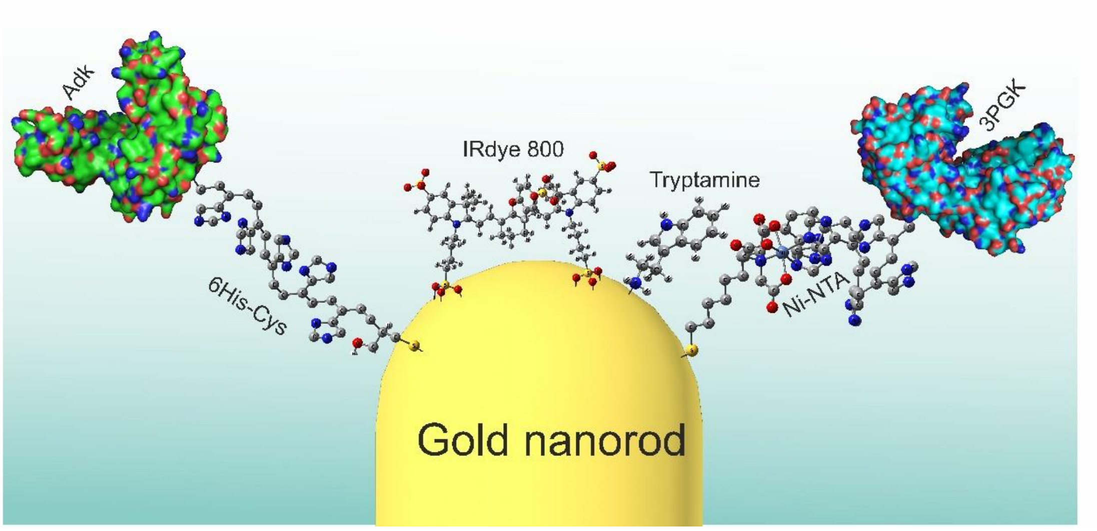


**Fig. S2.** Binding regime of all molecules tested. Alexa790 has no published structure so cannot be presented here, but linkages are proposed to be similar to those of IRDye800Cw.

**Protein Residue Components**

**Table S2 |** Residues present in designed Adk and 3PGK proteins.

| Adk | 3PGK |
|---|---|
| Ala (A) 10 | Ala (A) 49 |
| Arg (R) 12 | Arg (R) 20 |
| Asn (N) 3 | Asn (N) 13 |
| Asp (D) 11 | Asp (D) 31 |
| Cys (C) 1 | Cys (C) 1 |
| Gln (Q) 6 | Gln (Q) 2 |
| Glu (E) 23 | Glu (E) 33 |
| Gly (G) 18 | Gly (G) 35 |
| His (H) 10 | His (H) 20 |
| Ile (I) 16 | Ile (I) 21 |
| Leu (L) 18 | Leu (L) 38 |
| Lys (K) 22 | Lys (K) 32 |
| Met (M) 2 | Met (M) 9 |
| Phe (F) 6 | Phe (F) 15 |
| Pro (P) 15 | Pro (P) 15 |
| Ser (S) 4 | Ser (S) 11 |
| Thr (T) 6 | Thr (T) 13 |
| **Trp (W) 0** | **Trp (W) 2** |
| Tyr (Y) 7 | Tyr (Y) 8 |
| Val (V) 22 | Val (V) 38 |
| Pyl (O) 0 | Pyl (O) 0 |
| Sec (U) 0 | Sec (U) 0 |

**Evanescent intensity calculation**

We use a semi-empirical approach to evaluate the evanescent intensity $I$ of WGM near the nanorods. The field distribution $\mathbf{E}(\mathbf{r})$ of a specific WGM in microsphere can be numerically computed by using the formulas listed in ref.[3]. The effective mode volume of WGM is then derived as

$$V_{\text{eff}} = \frac{\int \epsilon(\mathbf{r})|\mathbf{E}(\mathbf{r})|^2 d\mathbf{r}}{\Lambda\epsilon(\mathbf{r}_0)|\mathbf{E}(\mathbf{r}_0)|^2}, \quad \textbf{[S1]}$$

where $\epsilon(\mathbf{r})$ denotes the spatial distribution of the relative permittivity and $\Lambda$ accounts for the local-intensity enhancement factor that arises from the localized surface plasmon resonance (LSPR) of the gold nanorod at the position $\mathbf{r}_0$. The typical value of $\Lambda$ in this work approximates 800. It should be noted that the definition of $V_{\text{eff}}$ here, i.e., Eq. [S1], is different from the one defined based on the maximum light intensity inside the microsphere.[3] An incident beam with the power $P$ pumps the WGM. At the steady state, the intracavity photon number reaches

$N_{\mathrm{in}} = \frac{\kappa_{\mathrm{in}}}{\kappa^2 \hbar\omega} P$. Here, $\kappa_{\mathrm{in}}$ and $\kappa$ are the coupling and total loss rates of the microsphere, respectively, $\hbar$ is the Plank's constant, $\omega = 2\pi c/\lambda$ is the angular frequency of the light, and $c$ is the speed of light. Thus, the light intensity at the position of nanorods is given by $I = \hbar\omega N_{\mathrm{in}} c/V_{\mathrm{eff}}$.

For each experimental data shown in figures, we measured the corresponding microsphere radius $R$, input power $P$, mode wavelength $\lambda$, total linewidth $\kappa$, and prism-microsphere coupling efficiency $S$. The effective mode volume $V_{\mathrm{eff}}$ is numerically computed based on $R$ and the prism-microsphere coupling rate is given by $\kappa_{\mathrm{in}} = (\kappa/2)\left(1 - \sqrt{1-S}\right)$. Then, the evanescent intensity $I$ can be evaluated accordingly. As an example, for IRDye800 we measured $R = 46.5$ μm, $P = 0.19$ mW, $\lambda = 780.029083$ nm, $S = 22\%$, and $\kappa = 503$ fm in experiment. $V_{\mathrm{eff}}$ and $\kappa_{\mathrm{in}}$ are respectively computed to be $V_{\mathrm{eff}} = 5.9 \times 10^{-18}$ cm$^3$ and $\kappa_{\mathrm{in}}/\kappa = 0.0584$ and then $I$ is evaluated to be $I = 36.2$ MW/cm$^2$.

**Theoretical model and fundamentals for single molecule absorption spectroscopy**

The resonance wavelength shift (from $\lambda$ to $\lambda'$) of WGM induced by the dielectric variation is expressed as[4]

$$\frac{\lambda' - \lambda}{\lambda} = \frac{\int [\epsilon'(\mathbf{r}) - \epsilon(\mathbf{r})] |\mathbf{E}(\mathbf{r})|^2 d\mathbf{r}}{2 \int \epsilon(\mathbf{r}) |\mathbf{E}(\mathbf{r})|^2 d\mathbf{r}}, \qquad \textbf{[S2]}$$

where $\epsilon(\mathbf{r})$ corresponds to the relative permittivity in the absence of the perturbation and $\epsilon'(\mathbf{r})$ denotes the relative permittivity in the presence of the perturbation. Specific to the experiment demonstrated, the resonance shift is caused by (i) the change of the relative permittivity at the location of ligand protein (approximately, the position of the gold nanorod $\mathbf{r}_0$) and (ii) the change of the relative permittivities of the environment (HEPES or Tris buffer) and microcavity due to the temperature rise (from $T$ to $T'$) caused by the fact that the protein heating water and microsphere. In addition, the LSPR effect has already been included in the relative permittivity $\epsilon'(\mathbf{r})$. Thus, the resonance shift $\Delta\lambda = \lambda' - \lambda$ may be re-written as (more detailed derivation can be found in Ref.[4])

$$2V_{\mathrm{eff}} \cdot \frac{\Delta\lambda}{\lambda} = \alpha_{\mathrm{ex}} + \sum_{i=\mathrm{w,s}} \frac{\epsilon_i(T') - \epsilon_i(T)}{\epsilon_i(T)} V_i, \qquad \textbf{[S3]}$$

with the excess polarizability of the protein $\alpha_{\mathrm{ex}} = \frac{\epsilon_{\mathrm{p}} - \epsilon_{\mathrm{w}}(T)}{\epsilon_{\mathrm{w}}(T)} V_{\mathrm{p}}$,[5] the relative permittivities of protein $\epsilon_{\mathrm{p}}$, water $\epsilon_{\mathrm{w}}(T)$, and microsphere $\epsilon_{\mathrm{s}}(T)$ at the temperature $T$, the protein volume $V_{\mathrm{p}}$, the effective volume of the heated water $V_{\mathrm{w}}$, and the effective volume of the heated microsphere $V_{\mathrm{s}}$. Note: the refractive index of proteins is in the order of $n = 1.4 - 1.5$, larger than that for water ($n = 1.33$).[6] This means $\alpha_{\mathrm{ex}}$ for all proteins used in this study (Adk, 3PGK) is always positive when measured in aqueous solution via the reactive sensing regime: the reactive regime response for protein binding will always be positive.

The relative changes of the water and microsphere permittivities caused by the temperature variation $\Delta T = T' - T$ are given by

$$\frac{\epsilon_i(T') - \epsilon_i(T)}{\epsilon_i(T)} = \frac{1}{\epsilon_i(T)} \frac{\partial \epsilon_i(T)}{\partial T} \Delta T = \frac{2}{n_i(T)} \frac{\partial n_i(T)}{\partial T} \Delta T, \qquad \textbf{[S4]}$$

with the refractive index $n_i(T)$ of water ($i = \mathrm{w}$) or microsphere ($i = s$) at temperature $T$. Equation [S3] is then re-expressed as

$$2V_{\text{eff}} \cdot \frac{\Delta\lambda}{\lambda} = \alpha_{\text{ex}} + \Delta T \cdot \sum_{i=\text{w,s}} \frac{2}{n_i(T)} \frac{\partial n_i(T)}{\partial T} V_i. \qquad \textbf{[S5]}$$

Since the LSPR only enhances the local field within a small region (hotspot) around the gold nanorod, both effective volumes $V_{\text{w,s}}$ of the heated water and microsphere are of the order of the hotspot volume ($\frac{2\pi}{3} \times (25\text{ nm})^3 = 3.3 \times 10^{-23}$ m$^3$). In addition, the change of refractive index of water with respect to the temperature $\left|\frac{\partial n_{\text{w}}(T)}{\partial T}\right| = -1.3 \times 10^{-4}$ K$^{-1}$ is negative and much larger than that of microsphere $\frac{\partial n_{\text{s}}(T)}{\partial T} = 1.3 \times 10^{-5}$ K$^{-1}$ and gives rise to the negative wavelength shifts.[7] Thus, the thermo-optic term associated with the microsphere in Eq. [S5] is negligible compared to that of water, and one obtains

$$2V_{\text{eff}} \cdot \frac{\Delta\lambda}{\lambda} = \alpha_{\text{ex}} + \frac{2V_{\text{w}}}{n_{\text{w}}(T)} \frac{\partial n_{\text{w}}(T)}{\partial T} \Delta T. \qquad \textbf{[S6]}$$

It is seen that the left side of the above equation is completely related to the microcavity (e.g., the effective mode volume and the resonance shift) and the right side of the above equation includes all environmental perturbations (i.e., the ligand–receptor interactions and the thermal effects). Since $V_{\text{w}}$ approximates the hotspot volume, we treat it as a water quasi-particle. Due to the negative value of $\frac{\partial n_{\text{w}}(T)}{\partial T}$, raising the local water temperature may result in a blue shift of the WGM resonance wavelength.

The heat absorbed by the protein per second is given by $h = \sigma_{\text{abs}} I$ with the absorption cross-section $\sigma_{\text{abs}}$ of the protein, and the light intensity $I$ at the position of the protein. It should be noted that the LSPR enhancement has been taken into account in $I$. Thus, the temperature increment $\Delta T$ is derived as

$$\Delta T = \frac{h}{\xi k_{\text{con}}} = \frac{\sigma_{\text{abs}} I}{\xi k_{\text{con}}}, \qquad \textbf{[S7]}$$

with the water's thermal conductivity $k_{\text{con}}$ and the effective heat transferring length $\xi$.[8] Considering the average of the resonance shifts, we arrive at Eq. [1].

**Table S3 | Examples of absorption cross-sections**

| Sample | 3PGK | Tryptamine |
|---|---|---|
| **Absorption cross-section by spectrometry at 780 nm, cm²** | 9.69 x10$^{-21}$ | 1.12 x10$^{-20}$ |
| **Absorption cross-section by TOP sensing at 780 nm, cm²** | 7.5 x10$^{-16}$ | 3.9 x10$^{-16}$ |

**Absorption Cross-Section Calculation**

The absorption cross-section $\sigma_{\text{abs}}$ was directly estimated with the following formula:

$$\sigma_{abs} = \frac{2.3 \cdot 10^3}{N_A} \varepsilon,$$

where $\varepsilon$ is the molar extinction coefficient.

The degree of labelling was calculated using equations supplied by ThermoFisher **(2)**, calculated as 0.65 and 0.76 for 3PGK-Alexa and Adk-Alexa, respectively.

$$A_{280nm}^{Alexa790} = A_{785} * CF_{280nm} \qquad CF_{280:785nm}^{Alexa790} = 0.08$$

$$A_{280nm}^{Protein} = A_{280nm} - A_{280nm}^{Alexa790}$$

$$[Protein] = \frac{A_{280nm}^{Protein}}{\varepsilon_{280nm}^{Protein} * L}$$

$$DoL = \frac{A_{785nm}}{[Protein] * L}$$

The extinction coefficients of Alexa790 and protein-Alexa790 conjugates were calculated by the below equation:

$$[Alexa790] = [Protein] * DoL$$

$$\frac{A_{280nm}^{Alexa790}}{[Alexa790]*L} = \varepsilon_{280nm}^{Alexa790}$$

$$\varepsilon_{280nm}^{Protein-Alexa790} = \varepsilon_{280nm}^{Alexa790} + \varepsilon_{280nm}^{Protein}$$

Where units of the values are:

Absorption (A): AU; Conversion Factor ($CF_{280:785nm}$): AU; Concentration ([x]): M; Pathlength (L): cm; Degree of Labelling (DoL): AU; Extinction coefficient (ε): $cm^{-1}M^{-1}$.

**Absorption spectra of subject proteins**

The absorption of 3PGK, 3PGK–Alexa, Adk, Adk–Alexa and tryptamine were evaluated by absorption spectroscopy using a Horiba Duetta absorption spectrometer (Fig. S3).

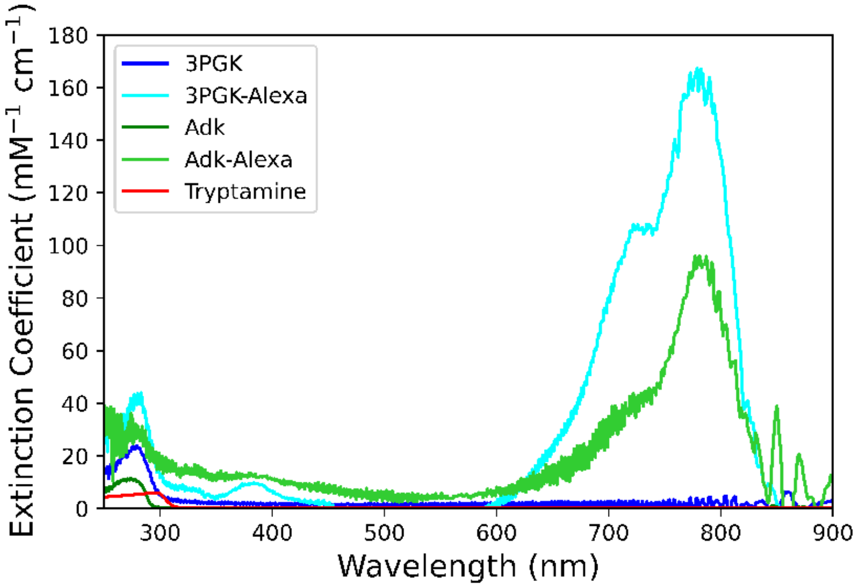


**Fig. S3.** Absorption spectra of the molecules under test.

**Temperature of the 'Hot Spot' Region**

We assessed the changes of temperature of nanorods when heated by radiation. Indeed, using the thermal diffusion equation, the surface temperature of a nanoparticle may be estimated with its absorption cross-section and size, incident power and thermal conductivity of the surrounding medium.[9,10] For gold nanoparticles used in an aqueous medium at WGM intensities of values we work with and typical values of absorption cross-sections of gold nanorods of ~$10^{-18}$ $m^2$ (taken from[11,12]), their temperature is changed within 20 K (Fig. S4).

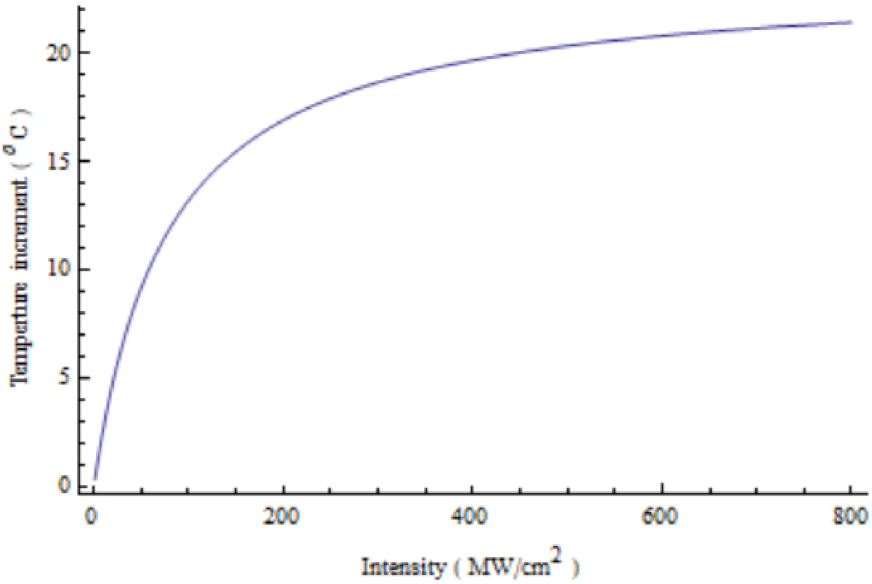


**Fig. S4.** Temperature of the locally heated area where molecules attach vs electric field intensity in this area.

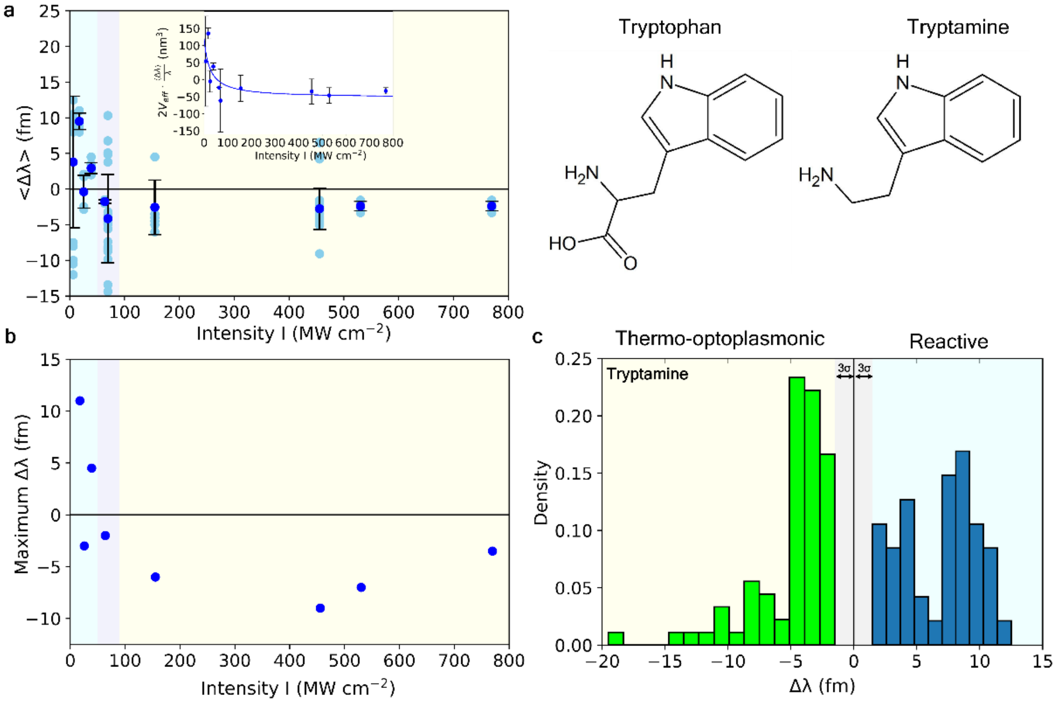

**Fig. S5. Single-molecule detection of small-molecule tryptamine. a,** Optoplasmonic sensing of single tryptamine molecules (blue circles). Averaged values of WGM wavelength shifts $\langle \Delta\lambda \rangle$ depend on the evanescent intensity $I$ of WGM around Au nanorods. Inset: Dependence of $2V_{\mathrm{eff}} \cdot \langle \Delta\lambda \rangle / \lambda$, with curve fit based on eq. (1) – $R^2$ = 0.5613. **b**, Maximal wavelength shifts at each intensity $I$ of WGM (by absolute values). Maximal wavelength shifts show binding at the tips of nanorods. **c,** Histogram of wavelength shifts over 117 individual datapoints. Both TOP sensing (green) and reactive mechanism (blue) are observed, dependent on evanescent intensity. Grey area indicates noise levels of triple the standard deviation, $3\sigma$. Tryptophan and Tryptamine structures are presented additionally.

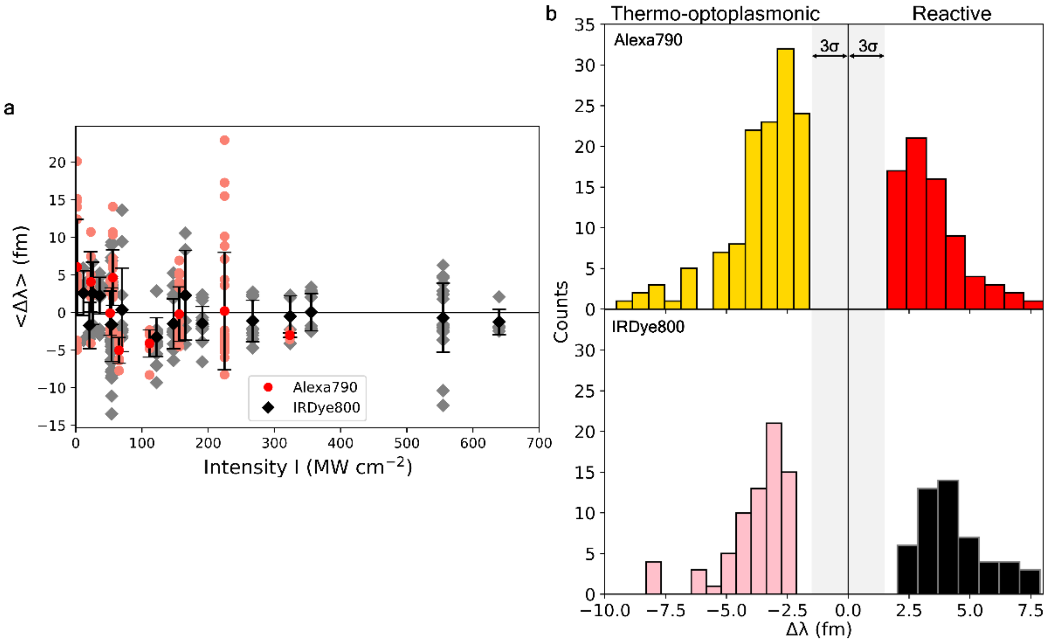


**Fig. S6. Single-molecule detection of Alexa and IRDye.** Resonant wavelength shifts and their averaged values. Alexa molecules (red circles) demonstrate sign-changing behaviour: red-shifted resonances at intensities up to 60 MW cm$^{-2}$ and blue-shifts at larger intensities. IRDye molecules (black diamonds) demonstrate blue-shifts even at smaller levels of intensities.

**Testing 3PGK molecules at higher intensities**

TOP sensing is limited by the intensity of WGM used for probing. At the current stage, it seems unnecessary to define maximum intensities allowed for experiments since they appear to be different for each molecule and does not provide additional information about absorption. Nevertheless, we performed experiments at higher intensities ($I$ = 625 MW cm$^{-2}$) that demonstrate anomalous large shifts (av. = 41.0 +/- 51.3) which are related to a different undescribed process: most likely protein aggregation (Fig. S7).

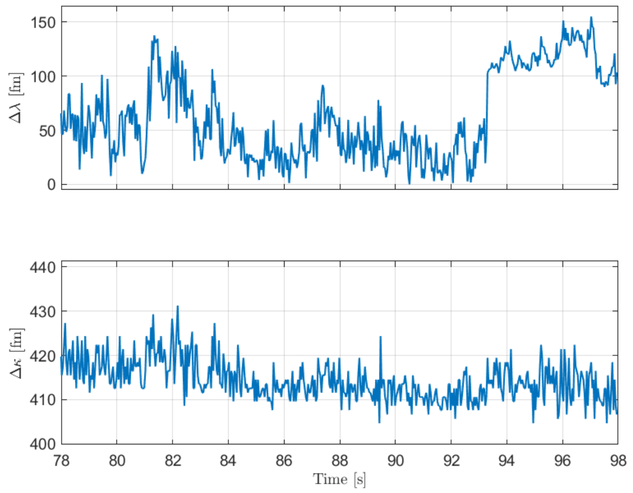


**Fig. S7.** Large wavelength shifts (Δλ) at *I* = 625 MW cm$^{-2}$. FWHM (Δκ) is also provided.
